# A robust image registration interface for large volume brain atlas

**DOI:** 10.1101/377044

**Authors:** Hong Ni, Chaozhen Tan, Zhao Feng, Shangbin Chen, Zoutao Zhang, Wenwei Li, Yue Guan, Hui Gong, Qingming Luo, Anan Li

## Abstract

Mapping the brain structures in three-dimensional accurately is critical for an in-depth understanding of the brain functions. By using the brain atlas as a hub, mapping detected datasets into a standard brain space enables efficiently use of various datasets. However, because of the heterogeneous and non-uniform characteristics of the brain structures at cellular level brought with the recently developed high-resolution whole-brain microscopes, traditional registration methods are difficult to apply to the robust mapping of various large volume datasets. Here, we proposed a robust Brain Spatial Mapping Interface (BrainsMapi) to address the registration of large volume datasets at cellular level by introducing the extract regional features of the anatomically invariant method and a strategy of parameter acquisition and large volume transformation. By performing validation on model data and biological images, BrainsMapi can not only achieve robust registration on sample tearing and streak image datasets, different individual and modality datasets accurately, but also are able to complete the registration of large volume dataset at cellular level which dataset size reaches 20 TB. Besides, it can also complete the registration of historical vectorized dataset. BrainsMapi would facilitate the comparison, reuse and integration of a variety of brain datasets.

## Introduction

Mapping brain structures in three-dimensional is necessary to thoroughly understand of the brain functions (Huang and Luo, 2015). Creating a comprehensive space to map various brains will encompasses complex spatiotemporal information that can greatly facilitate the comparison (Badea et al., 2007), reuse (Boline et al., 2008) and integration (Gupta et al., 2000) of brain datasets. Drawing a stereotaxic brain atlas (Dong, 2008; Goldowitz, 2010) provides a unified spatial reference for addressing this issue. However, with the rapid development of high-resolution whole-brain microscopic imaging (Gong et al., 2016; Li et al., 2010; Ragan et al., 2012), the obvious heterogeneous and non-uniform (Lawson et al., 1990) characteristics of brain structures at cellular level make it difficult to map various experimental datasets from different individuals, modalities to a standard anatomical coordinate space by using uniform registration methods (Klein et al., 2009), which used in previous macroscopic level datasets such as magnetic resonance imaging (MRI). In addition, with the produced large volume dataset during imaging, we urgently need a robust nonlinear registration pipeline that can register massive spatial information datasets, and accurately position at the cellular level.

Previous studies (Kakadiaris et al., 2004; Maintz and Viergever, 1998; Ohnishi et al., 2016; Zitova and Flusser, 2003) have been conducted to solve the registration of three-dimensional whole-brain datasets, especially within the MRI field. Among these studies, gray-level based registration algorithms (Klein et al., 2009) can effectively achieve the nonlinear registration of MRI datasets with uniform signals. These methods can also be applied to optical microscopic images, e.g. Leonard et al. (Kuan et al., 2015) processed the serial two-photon (STP) datasets to obtain an average brain. However, the optical microscopy images are susceptible to sample preparation and imaging processes, which will lead to the calculation of energy function of gray-level based method falling into the local minimum. For more complex optical microscopy images, feature-based registration algorithms (Fürth et al., 2018; Sergejeva et al., 2015; Zitova and Flusser, 2003) are the better choice because the factors that affect registration accuracy can be changed from grayscale to the extracted feature information, and then the bottle neck evolves to how to accurately and objectively extract sufficient feature points. Ohnishi et al. (2016) completed the registration of two-dimensional microscopy images to MRI data by extracting manually feature points, but its number was limited and subjective. Wang et al. (2014) used automatic methods to extract feature points and achieved the registration of histopathological data, but the accuracy of automatic recognition was greatly affected by data quality. Fürth et al. (2018) combined the advantages of artificial and automatic methods and achieved the registration of two-dimensional continuous microscopy images to a brain atlas. Briefly, when neuroscientists are struggling to obtain a valuable experimental dataset, they often find that the existing registration method is not very effective because of excessive reliance on image quality or experience.

Another challenge is the TB-scale large volume whole-brain datasets brought by imaging at cellular level. Generally, registration tools widely used in the biomedical imaging field, such as ITK (Ibanez et al., 2005) can achieve nonlinear registration only at GB level, the maximum amount of data reported in the literature (Niedworok et al., 2016) is approximately 1 GB, specifically a 12.5 μm^3^ resolution whole-brain dataset. Clearly, the current nonlinear registration algorithm is a global optimization solution that requires a large amount of memory consumption and iterative computation and is not suitable for block parallelism. Therefore, achieving robust nonlinear registration at cellular level with a TB-scale large volume dataset is difficult.

Here, we proposed BrainsMapi, a robust registration interface to accurately map three-dimensional brain image datasets to the standard brain atlas at cellular level. It can be applied to various datasets with sample tearing and streak image datasets, different individual and modality datasets, and we have registered four types of datasets (MRI (Johnson et al., 2010), Nissl staining (Micro-Optical Sectioning Tomography, MOST) (Li et al., 2010), propidium iodide (PI) staining (Brain-wide Precision Imaging system, BPS) (Gong et al., 2016) and STP (Ragan et al., 2012)) to the Allen Common Coordinate Framework version 3 (Allen CCFv3) for a robust demonstration (Goldowitz, 2010; Kuan et al., 2015). The registration results were highly accurate at the brain region level (Dice score > 0.9), and there was no significant difference with manually results in the identifiable nuclei level at 10 μm resolution (P > 0.05). BrainsMapi can also process large volume three-dimensional brain image datasets, and we demonstrated the nonlinear registration of whole brain dataset with more than 10 tera voxels. In addition, It is compatible with historical data and are able to register existing labeled data, which can integrate datasets comprising neuroscience information from different experimental brains into a common brain space.

## Materials

### Model data

To visually demonstrate the effectiveness and robustness of BrainsMapi, we designed five simple, cartoon models with smiley and crying faces. The process is as follows.
1. Fixed model: A round smiley face in which the eyebrows, eyes, and mouth were marked with different gray level values.
2. Model1: Manually designed a deformation, deformed the fixed model, and made it a crying face.
3. Model2: Using the histograms of corpus callosum (cc), hippocampal region (HIP) and Cerebellum (CB) of the Allen CCFv3 Nissl stained dataset, randomly assigned the eyebrows, eyes, and mouth of Model1, respectively.
4. Model3: Added streak noise to Model2 horizontally and vertically by using a sine function.
5. Model4: On the basis of Model3, set a gray level of zero in a triangular area of the mouth to simulate a tearing condition for the samples.

### Biological datasets

We employed 6 whole-brain datasets and 1 metadata to validate BrainsMapi. All of the animal experiments followed procedures approved by the Institutional Animal Ethics Committee of Huazhong University of Science and Technology.

Dataset 1 is from the CCFv3 of Allen Institute. This dataset is the average brain obtained by continuously averaging the 1,675 STP image datasets. The downloads of the average brain, annotation file, and individual Nissl dataset at four resolution levels of 100 μm, 50 μm, 25 μm, and 10 μm are open available at http://brain-map.org. Here we chose the average brain dataset of 10 μm resolution.

Dataset 2 is the image volumes representing the canonical Waxholm Space (WHS) adult C57BL/6J mouse brain. Five datasets are provided, and three of the datasets were acquired by Duke Center for in-vivo microscopy using three MRI imaging models T1, T2, and T2*, one is the Nissl stained dataset obtained by Drexel University, and one is the manually Labeled Atlas dataset. The datasets are available at https://www.nitrc.org/. Here we chose the T2*-weight MRI image dataset.

Dataset 3 is the whole-brain image dataset of a Nissl stained C57BL/6 adult mouse imaged by MOST. The three-dimensional dataset with 1 μm axial resolution and 0.35 μm horizontal resolution was acquired in 7 days using the MOST automatic slicing imaging system. The coronal slice number and original data size were approximately 11,000 and 4 TB respectively.

Dataset 4 is from the whole-brain dataset of dual-color labeled Thy1-GFP M-line transgenic mice imaged by BPS. The co-localized fluorescent-labeled neurons and counterstained cell bodies dataset in brain wide with 2 μm axial resolution and 0.32 μm horizontal resolution was acquired. The coronal slice number and original data size of one channel dataset were approximately 4,800 and 3 TB respectively. Here we chose the cytoarchitectonic channel dataset. In addition, we identified and reconstructed the barrel cortex neurons of the GFP channel. Approximately, a total of 40 neurons were used as Metadata 1 (Gong et al., 2016).

Dataset 5 is the specifically selected problematic dataset, which contains streaks caused by uneven staining and illumination and sample tearing phenomenon. This dataset will be used for comparison of later results. The imaging system and staining method are the same as those obtained by Dataset 4. The only difference is that the axial resolution is 1 μm.

Dataset 6 is a collection of published article (Niedworok et al., 2016) from the Division of Neurophysiology, MRC National Institute for Medical Research, London. Briefly, it is a dataset of autofluorescent C57BL/6 mouse imaged by STP microscopy. The original dataset has an axial resolution of 5 μm and a coronal resolution of 0.32 μm. The dataset is available at http://www.swc.ucl.ac.uk/aMAP.

Please refer to the SI Table 1 for the main information of all image datasets (SI Figure 1) and metadata.

**Figure 1.**
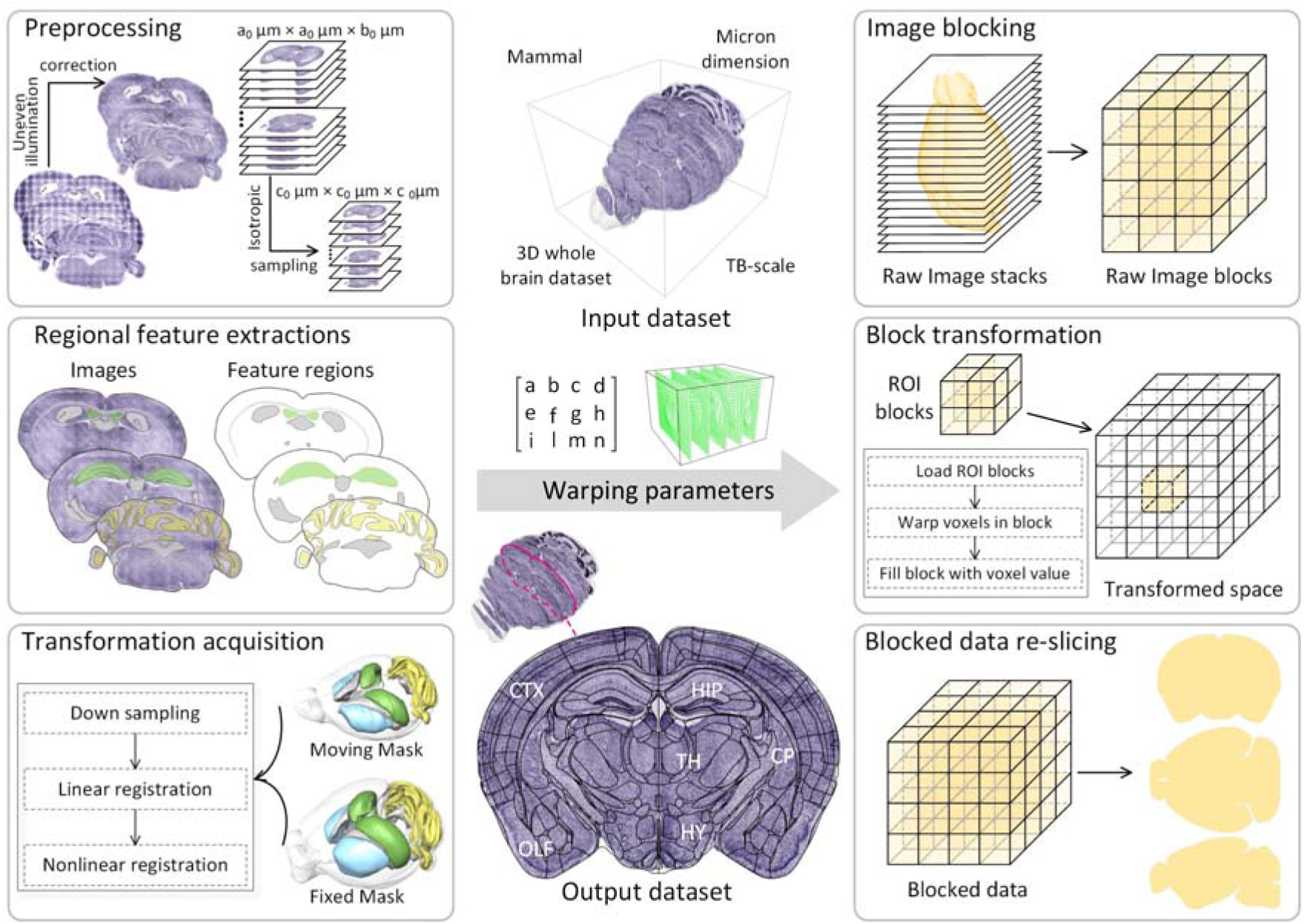
Registration pipeline for large volume brain datasets. Input dataset: mammal, three-dimensional whole brain, micron dimension, TB-scale dataset. The crucial steps for ensuring the robustness of registration, including preprocessing, regional feature extractions, and transformation acquisition are on the left. The obtained warping parameters are input to the right, and the registration for large volume brain datasets, including image blocking, block transformation and blocked data re-slicing. The registered result is obtained in three anatomical sections.

### Computing Environments

In this paper, we used two computing devices. A graphical workstation was equipped with 20 cores (Intel Xeon E5-2687w×2), and 128 GB of RAM. Another device is HPC Cluster with 20 nodes. Each node was equipped with 20 cores (Intel Xeon E5-2660 V3×2) and 128 GB of RAM and connected with the Lustre file system via a 10 Gb Ethernet.

## Methods

The whole pipeline of BrainsMapi is shown in Figure 1. We proposed a regional feature extraction method that can accurately, objectively and sufficiently extract features. This method would not be affected by factors such as sample defect, image quality and imaging modality that are crucial for ensuring the robustness of registration. Based on these, with a parameter acquisition and high-resolution transformation strategy we achieve the whole-brain nonlinear registration of the TB scale dataset at single-cell resolution. We blocked a large volume dataset, and used high-performance compute to efficiently transform each block in parallel. The pipeline included image preprocessing, regional features extractions, accurate transformation acquisition and nonlinear transformation for a large volume dataset. We will describe the technical detail below.

### Image preprocessing

Image preprocessing aims to obtain high quality images for good registration results (Figure 1 Preprocessing). First, denoising the original image, such as brightness correction and light-field correction to reduce the uneven staining and illumination of optical microscopy imaging (Ding et al., 2013) was necessary, next, we performed a preliminary rotation on the dataset to ensure proper orientation with the reference image. Then the corrected dataset was sampled isotopically. Finally, we used the adaptive threshold method (Otsu, 1979) and morphological operations (hole filing, opening and closing operators) to extract the mouse outline and remove the background.

### Regional feature extractions

The accuracy of feature selection has a direct relationship with the registration results. Here, using the advantages of brain anatomy information, we proposed a method to extract anatomically invariant regional features to register two brain datasets. The extracted regional features with anatomical meanings, that is, brain regions or nuclei that were also conservative to delineate the boundaries of conservative brain regions or nuclei in three-dimensional space. Compared to observing a single anatomical point in three-dimensional brain space, extracting the boundaries of anatomical regions is more accurate, objective, and contains information such as shape and size.

Here, an interactive segmentation tool, Amira (version 6.1.1; FEI, Mérignac Cedex, France), was chosen to perform the feature extraction procedure (Figure 1 Extraction of regional features). Briefly, the selection of extracted brain regions and nucleus needs to follow these three criteria.
1. Distributed throughout the brain to ensure the accuracy of registration in brain-wide.
2. Easily identified anatomical features to ensure accuracy and objectiveness for feature extraction.
3. Selection of conservative brain regions or nuclei that are guaranteed to occur in every brain as the basis of registration to ensure the correctness of the registration.

Based on the criteria, we selected following features in the whole-brain with the guidance of anatomists: Outline, anterior commissure, olfactory limb (aco)/ anterior commissure, temporal limb (act), CB, cc, caudoputamen (CP), fasciculus retroflexus (fr), HIP, medial habenula (MH), facial nerve (VIIn), mammillothalamic tract (mtt), paraventricular hypothalamic nucleus (PVH), pontine gray (PG), lateral ventricle (VL), and fourth ventricle (V4). We have shown these regions in SI Figure 3. The entire regional feature extraction step was performed at 10 μm resolution, and all features were saved in the 3D-TIF image file.

### Accurate transformation acquisition

After accurately extracting features brain wide, we needed to map these features to acquire accurate transformation parameters (Figure 1 Transformation acquisition). The process of transformation acquisition is an optimal problem. The moving image is warped to the fixed image by initial transformation parameters, and the similarity metric of moving and fixed images is used as an energy function. Iteratively, the transformation parameters are updated to achieve an optimal solution and obtain the corresponding transformation. The transformation is composed of linear and nonlinear parameters, where the linear parameter can be simply represented by matrix *M* describing the translation, rotation, and scaling of 12 degrees of freedom, and the displacement field *φ* can represent the nonlinear parameter. Here, we chose a recent nonlinear registration method, Symmetric Diffeomorphic Normalization (SyN) (Avants et al., 2008). SyN customizes the symmetry deformation based on the standard method of the Large Deformation Diffeomorphic Metric Matching (LDDMM) proposed by Beg (Beg et al., 2005). SyN can flexibly record the displacement for each pixel with a large deformation and produces a diffeomorphic transformation of symmetric and invertible. This symmetrical method of processing direct and inverse simultaneously is also reflected in its energy function (Equation 1).

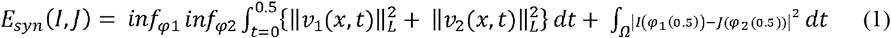

Where *φ*1 and *φ*2 are the diffeomorphism field in opposite directions of domain *Ω* indexed by time *t, t* ∈ [0,1], and *v*1 and *v*2 are the velocity field in opposite directions. Physically, the distance drives each pixel to move, which is determined based on the image potential energy. When the original images are replaced by our extracted regional features, the movement of each pixel is determined by the potential energy of features, which will not be disturbed by large differences in image gray level and will not fall into local minimums caused by image noise. In short, when we replace I and J which represent the original images with the feature images *I*_0_ and *J*_0_, we can obtain accurate transformations.

We used the five-pyramid strategy in both linear and nonlinear registration for acceleration, and mutual information was used as the similarity measure. The entire transformation acquisition step was approximately 3 hours. We obtained the linear matrix M, the nonlinear direct displacement field *φ*1 and the inverse field *φ*2.

Furthermore, the displacement was presented in a grid form to intuitively illustrate the nonlinear deformation effect of the diffeomorphism method based on regional features (Figure 2). Three-dimensional nonlinear displacement (Figure 2B) and nonlinear registration results (Figure 2C) were obtained by the nonlinear registration on the linear results (Figure 2A). The global and local nonlinear deformation effects where be observed from the coronal, sagittal and horizontal sections respectively (Figure 2D). For comparison, we also show the registration effects with the atlas line superimposed the nonlinear results (Figure 2E).

**Figure 2.**
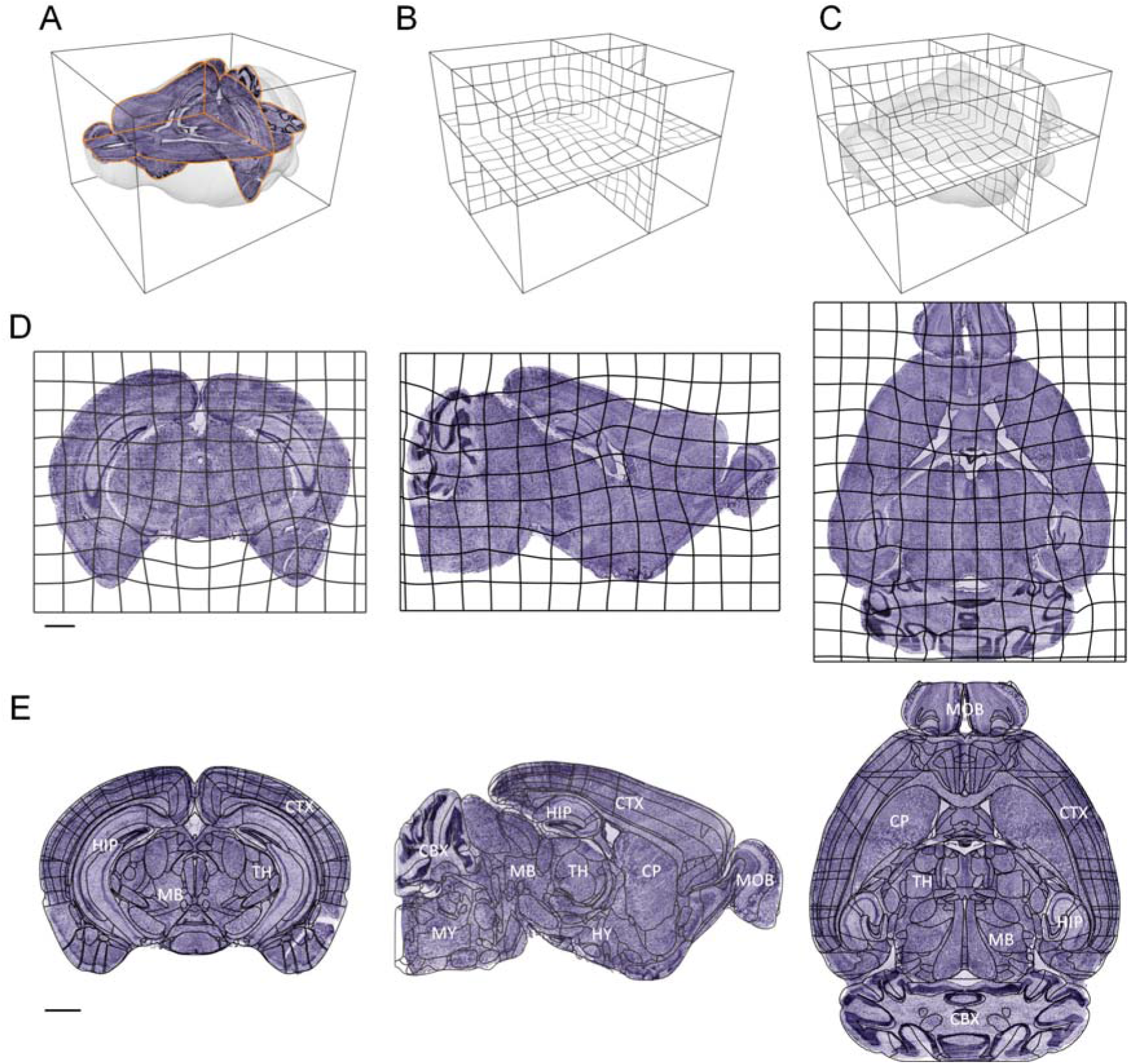
Application of nonlinear deformation field. (A) The three-dimensional rendering of the brain outline and three anatomical sections (coronal, horizontal and sagittal) before nonlinear registration. (B) Three-dimensional displacement. (C) Three-dimensional deformation field applied to the original three-dimensional dataset. (D) A two-dimensional grid shows the application of deformation fields in coronal, sagittal, and horizontal planes. (E) The registration of coronal, sagittal and horizontal sections correspond to (B), respectively. Scale bars: 1 mm.

### Nonlinear transformation for a large volume dataset

The existing nonlinear registration algorithms load the entire volume into memory for calculation. However, this strategy will dramatically increase the memory consumption and running time for the TB-scale dataset, simply stacking hardware cannot solve the problem of nonlinear registration of TB-scale dataset. Here, a parameter acquisition and high-resolution transformation strategy were proposed to achieve nonlinear registration of the TB scale dataset at single-cell resolution.

First, the transformation parameters for the low-resolution dataset (10 μm isotropic) were obtained based on the above transformation strategy. Then we acquired the transformation parameters for the high-resolution dataset as follows:

1. For linear parameters, the linear matrix *M* (low-resolution) was multiplied with a scale matrix to produce the high-resolution linear matrix (*M*’);
2. Nonlinear parameters are represented by deformation fields, which express the displacement for each voxel in three-dimensional space in the form of three channels *φ_x_, φ_y_, φ_z_*. By simply up sampling the low-resolution displacement will generate a TB-scale high-resolution displacement, which encounters storage and computation problems. Conversely, we directly calculated each voxel displacement (*φ_x_ (x, y, z), φ_y_ (x, y, z), φ_z_ (x, y, z)*) by interpolation during the transformation process according to low resolution displacement fields.

*P(x, y, z*) dentes the space coordinate of voxel P before registration, and the coordinates after linear and nonlinear registration are *P’ (x’, y’, z’*) and *P”(x”, y’’, z’’*), respectively. The mapping relationship is shown in Equation 2.

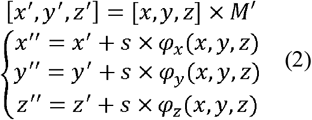

Based on the above mapping relationship, we established a transformation method for large volume datasets. Raw image sequences were partitioned into many cubes using TDat tools (Li et al., 2017b) (Figure 1 Image blocking). The transformed space was precalculated according to the size of fixed images and scaling factor. Next, we applied transformation parameters for each block in transformed space separately. We loaded only a small range of data into memory to solve the contradiction of the TB-scale dataset and the limitation of memory. The details are as follows (Figure 1 Block transformation).
1. For each block in transformed space, we obtained the corresponding blocks in their original space by transforming all points on six surfaces of the block with the mapping relationship (Equation 2). Then, the identified blocks (ROI blocks) in original space were loaded to memory.
2. For every voxel in a block, we calculated the spatial coordinate of the corresponding voxel in ROI blocks by the mapping relationship (Equation 2) and used the grayscale value of the coordinate as this voxel.
3. Each block was written to a disk in three-dimensional image format with gray values.

After the process above, we acquired the registered three-dimensional image dataset at original resolution, which was in the TDat format. Finally, the registered two-dimensional image sequences of three anatomical sections (coronal, horizontal, sagittal) were generated by re-slicing (Figure 1 Blocked data re-slicing).

During the process, parallel technologies of process-level message passing interface (MPI) and thread-level OpenMP were applied in data reading, writing and calculating operations in a multicomputer environment to efficiently achieve the nonlinear registration of the TB-scale whole-brain dataset.

### Vectorized dataset registration

Integrating a dataset such as cells (Peng et al., 2017; Zhang et al., 2017), neurons (Gong et al., 2016; Li et al., 2017a), and vessels (Xiong et al., 2017) requires mapping them into a standard space. These vectorized datasets are constructed of point set and coordinates. Using neurons as an example, traced neurons were saved as SWC files in the form of sequence points. In the same manner as the image dataset transformation, we transformed each point, and realized the registration of these vectorized datasets.

### Multilevel quantitative evaluation

Most evaluation methods (Christensen et al., 2006; Kim et al., 2015; Klein et al., 2009; Oh et al., 2014) use feature points or manual segmentation. According to the literature (Zitova and Flusser, 2003), the most effective way is subjective judgment by anatomists. We think that simply selecting a few points cannot completely assess the registration results, and the segmentation results of individuals are also subjective. Here, we designed a set of multilevel quantitative assessment methods, from coarse to fine and assessed the registration results at the brain-region level to the identifiable nuclei level at 10 μm resolution accurately and objectively.

By design, we first assessed the accuracy of the brain-region level, a coarse assessment. A total of ten brain regions of interest were chosen throughout the brain: Outline, CB, CP, hindbrain (HB), HIP, hypothalamus (HY), isocortex (ISO), midbrain (MB), pons (P) and thalamus (TH). We manually segmented the moving image after registration, and the segmentation results were regarded as a silver standard (Klein et al., 2009) instead of a golden standard. To make the segmentation results more accurate and reduce the workload, we did not segment each complete brain region slice by slice, because of the extreme indistinguishability of the start and end of the brain regions. The brain regions were middle intercepted and 50 image sequences were selected at equal intervals and segmented slice by slice. All manual segmentation results were referenced to the Allen CCFv3. Finally, the dice score (Equation 3) was selected as the evaluation measure.

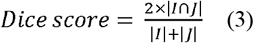

*I* is the moving image after registration, *J* is the fixed image, and ∩ denotes the intersection of two images. The dice score was calculated in two-dimensional, the number of dice scores for each brain region was 50.

In addition to the design of identifiable nuclei level evaluation at 10 μm resolution, an aMAP (Niedworok et al., 2016) approach was used as a reference. Nine nucleus were selected brain-wide (anterior cingulate area (ACA), primary visual area (VISp), primary somatosensory area (SSP), reticular nucleus of the thalamus (RT), ventromedial hypothalamic nucleus (VMH), periaqueductal gray (PAG), subiculum (SUB), entorhinal area, lateral part (ENT1), medial vestibular nucleus (MV)), and included both distinguishable and indistinguishable boundaries. Twenty-two trained technicians segmented these regions in five registered datasets (four type). First, we selected a representative coronal section in the CCFv3 for each nuclei, and a stack (40 sequences) of corresponding position in the registered datasets was also provided. Then, these 22 individuals needed to identify a single coronal section from the stack that they considered most similar to the reference coronal section and segmented it. Based on this procedure, the STAPLE algorithm (Warfield et al., 2004) was used to fuse 22 human segmentation results, and obtain the SPATLE results. The dice scores of each segmentation result and SPATLE results were calculated as human performance (HP), and the dice scores of the CCFv3 and SPATLE results were calculated as registration performance (RP). Then, correlation analysis was performed.

## Results

### A model experiment for robust demonstration

Using the corresponding constructed models according to the complex situations during brain image registration such as sample deformation, weak signal-to-noise ratio (SNR), streak noise and sample tear, we demonstrated the robustness of our proposed BrainsMapi and other registration methods in response to various complex situations.

We simulated sample deformation in Model1 and added a non-uniform gray assignment to simulate non-uniform signal or weak signals (eyebrows) in Model2. Model3 was added worse streak noise. On the basis of above steps, we added a sample tear situation to Model4 (Figure 3 and Materials). When the crying face turns into a smiling face, it proves that good registration effects have been achieved. As shown in the last column in Figure 3, regardless of deformation, weak SNR, streak noise and tearing, proposed method could obtain good registration results.

**Figure 3.**
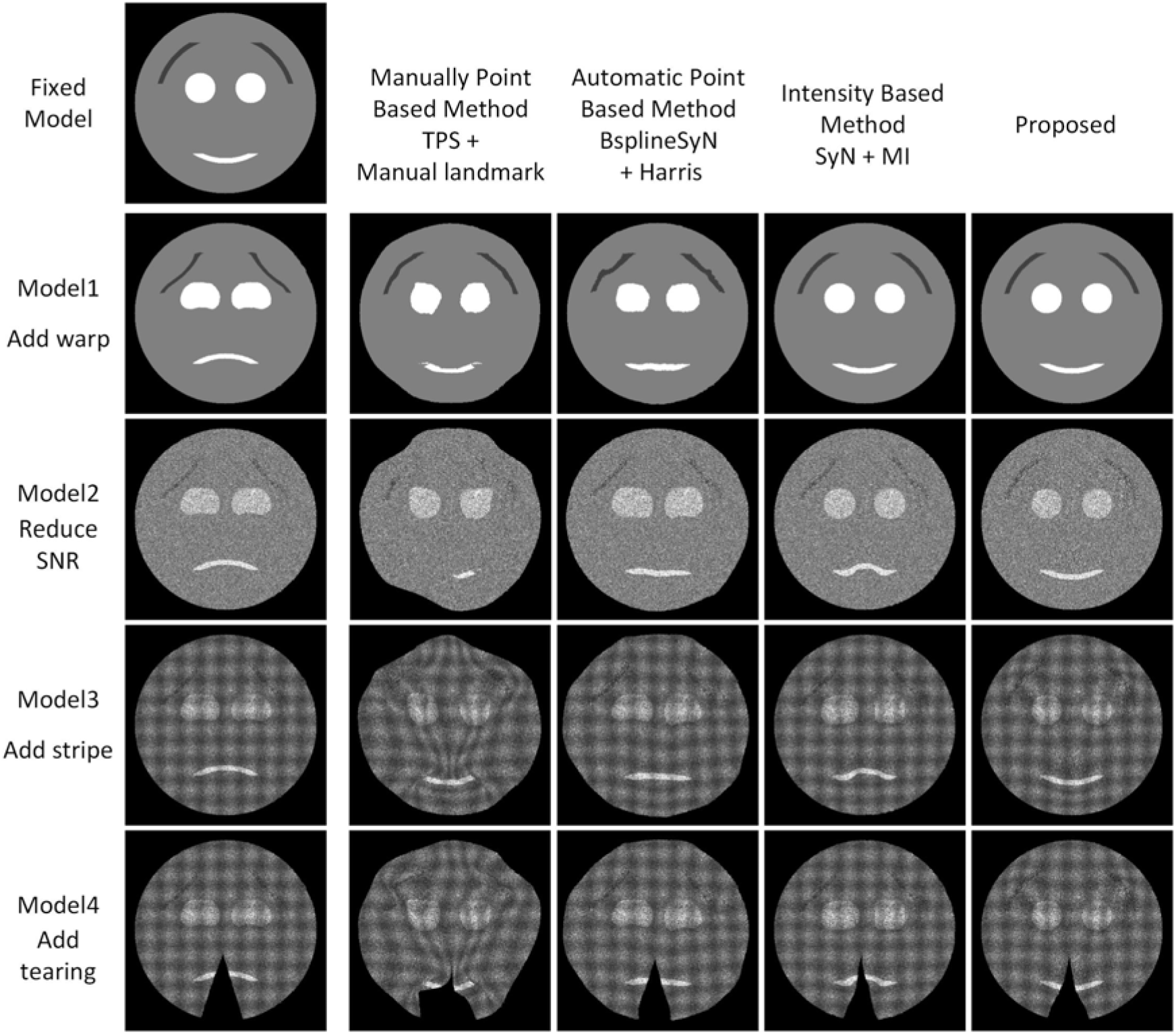
Registration effects for model data. The comparison of three traditional registration methods with our proposed method. The first row of smiles is the fixed model. Rows 2 to 5 correspond to the registration results of three traditional registration and proposed BrainsMapi in different modeled situations.

In contrast, we also listed the effects of other registration methods on the model data. The results of manually selecting feature points (eg. TPS with manually landmark) are randomness due to the subjectivity, limited numbers and inconsistent location of the selected feature points (Figure 3, second column). Accurate automatic extraction of feature points (eg. BSplineSyN with Harris) is difficult in weak SNR and strong noise images, resulting in inaccurate registration (Figure 3, third column). The gray-level based registration method (eg. SyN with MI) also cannot be applied to weak SNR, strong noise interference and tearing images (Figure 3, fourth column). Three-dimensional model datasets registration results are also provided in supplementary materials (SI Figure 5).

**Figure 4.**
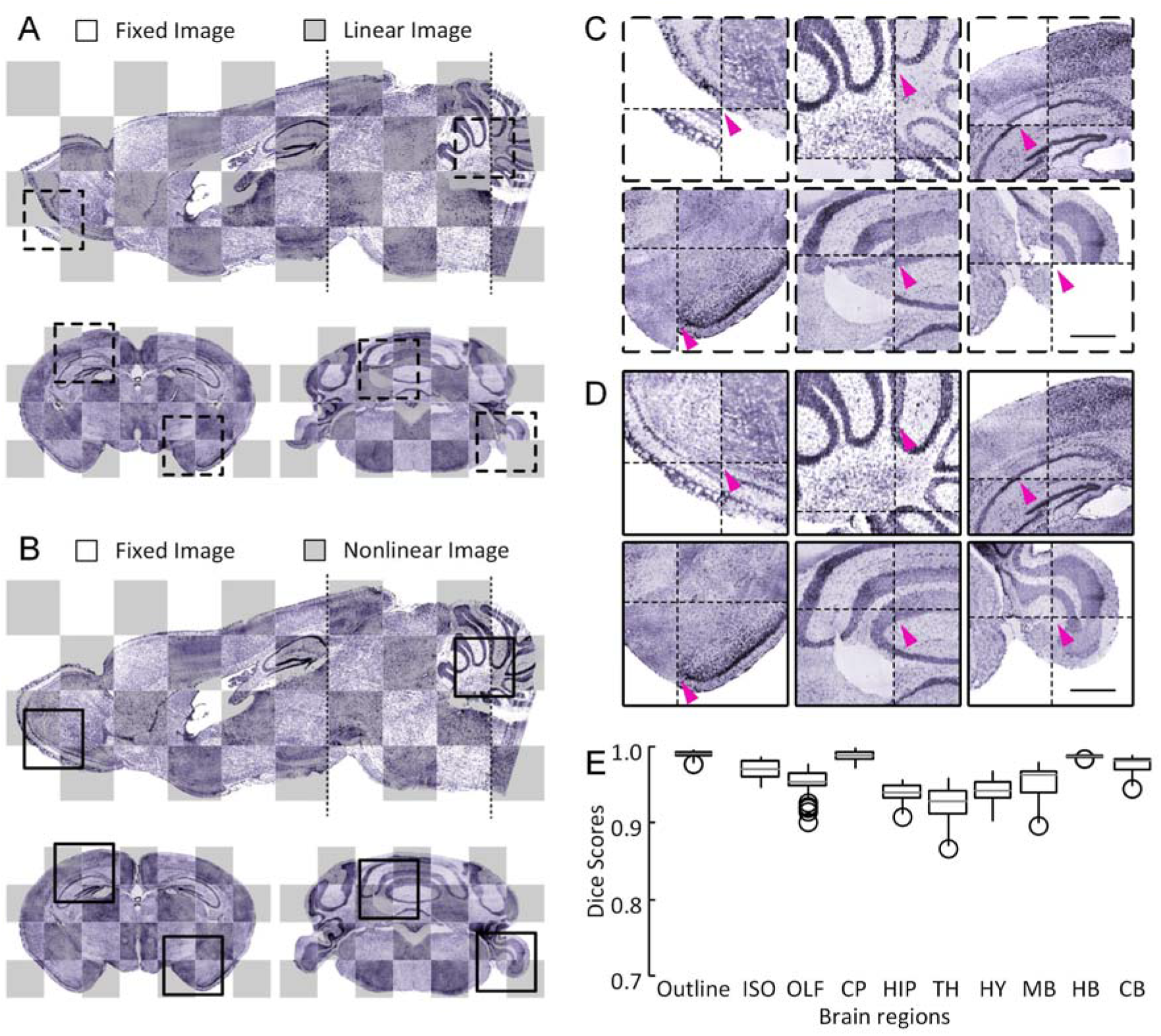
Registration effects for sample tearing and streak images. (A) The checkboard reconstructions of linear registration of one sagittal and two coronal slices indicated by dashed lines in the sagittal slice. The projection thickness is 10 μm. (B) The checkboard reconstructions of nonlinear registration results of one sagittal and two coronal slices indicated with dotted lines in the sagittal slice. The projection thickness is 10 μm. (C) The enlarged views of linear results of local regions indicated with corresponding dotted box in (A). (D) The enlarged views of nonlinear results of local regions indicated with corresponding dotted box in (B). The thin cross dotted line in (C, D) represents the grid of the checkboard. (E) A quantitative evaluation at the brain-region level.

**Figure 5.**
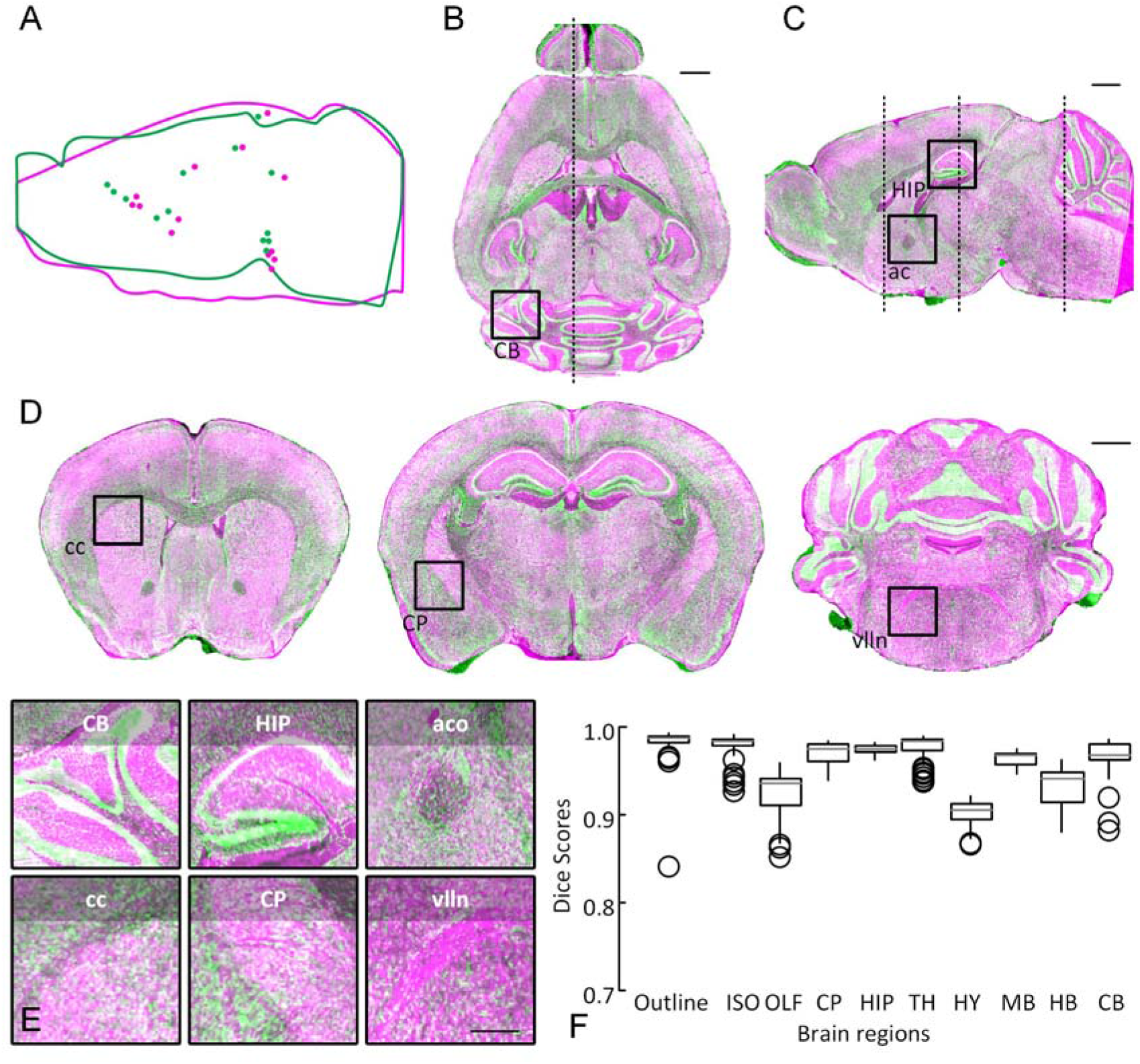
Registration effects for large different individuals. (A) The schematic diagram of thirteen pairs of landmarks and brain outlines, fixed image (purple), moving image (green). (B) A horizontal reconstruction at the location indicated by a dot outline in (A), merge the fixed image (purple) with the registered moving image (green). The projection thickness is 10 μm. (C) A sagittal reconstruction at the location indicated by a dotted line in (B), merge the fixed image (purple) with the registered moving image (green). The projection thickness is 10 μm. (D) The 10 μm thick projection of several merged coronal sections indicated by dotted line in (C). (E) Merged image of enlarged views of local regions indicated with text annotations and corresponding boxes in (B, C, D). (F) A quantitative evaluation at the brain-region level. Scale bars: (B, C, D) about 1 mm, E 0.5 mm. Cerebellum (CB), Hippocampal region (HIP), anterior commissure, olfactory limb (aco), corpus callosum (cc), caudoputamen (CP), facial nerve (VIIn).

### Registration for sample tearing and streak image dataset

The complications simulated in the model data also existed in brain images. There are large differences in image quality even with the same modality images. These images are easily disturbed in the processes of sample preparation and imaging, such as obvious sample tearing and streaks caused by uneven illumination (SI Figure 2). Gray-level or feature based registration methods cannot easily to solve these problem, which prevents us from comparing and analyzing intro-modal datasets.

Using Dataset 4 as a reference brain, we presented an intra-modal registration for poor quality (sample tearing and streaks) images by aligning Dataset 5 to the reference brain based on BrainsMapi. We presented the registration results in the form of checkerboards in Figure 4. The global orientation of the mouse brain was corrected after linear registration (Figure 4A), and the local brain regions and nucleus were adjusted after nonlinear registration (Figure 4B). Compared to linear results (Figure 4C), nonlinear registration resulted in good local correction of regions such as olfactory areas (OLF), CB, HIP, and paraflocculus (PFL) (Figure 4D). The small purple arrows in Figure 4CD indicate that the misalignment positions were corrected after nonlinear registration.

In addition, we evaluated the registration accuracy at brain-region levels (See method). In the box plot (Figure 4E), the median dice score for all brain regions was between 0.9 and 0.99. Generally, a dice score is above 0.8 indicates that good registration effects have been achieved (Klein et al., 2009; Niedworok et al., 2016).

### Registration for large different individuals

In addition to the differences in image quality, the more general differences in the experiment are individuals. Here, we expect to show registration results of large different individuals. Using a way of measuring the distance of anatomical landmarks (Kim et al., 2015), we found that the greatest different individual pair was Dataset 2 and Dataset 4 between the four dataset types (Dataset 2 (MRI), Dataset 3 (MOST), Dataset 4 (BPS) and Dataset 6 (STP)) (SI Figure 4), and we also shown the distance between these two datasets in Figure 5A (median 596.9 μm).

Using Dataset 2 as a reference brain, Dataset 4 was registered. By merging Dataset 4 (green channel) to Dataset 2 (purple channel), we presented the inter-modal registration results of large individual differences with horizontal (Figure 5B), sagittal (Figure 5C) and several coronal sections (Figure 5D). Roughly, the brain outline and big brain regions could be well aligned. Moreover, the corresponding enlarged views were given in Figure 5E, nucleus such as CB, HIP, aco, cc, CP, VIIn could also be well mapped.

Here, we also conducted a quantitative assessment of brain-region levels. In the box plot (Figure 5F), the median dice score for all brain regions was between 0.9 and 0.99.

### Register to a reference atlas for multi-modal registration

We presented a special registration type in this section that was registered to the reference atlas. Using direct or inverse registration to the reference brain space, we were able to acquire the spatial information of the original dataset, which is the key to integrating multi-modal image dataset into a common brain space.

We registered four types of datasets (Dataset 2-4 and 6) to Allen CCFv3 (Dataset 1) and presented the registration results. From top to bottom, the information corresponds to the registration results (Figure 6) of Dataset 2-4 and 6. We selected three coronal sections in the whole-brain with the form of the Allen CCFv3 on the left and the yellow dotted line of the Allen CCFv3 superimposed on the original image to show our results. The brain outline and big brain regions such as HIP and CB were well aligned. Brain regions without obvious anatomical boundaries such as TH, HY, MB and CB also had a good alignment with the Allen reference which could be judged from their spatial orientation. Moreover, we presented the enlarged views of local regions of nucleus for each data type. Nucleus with distinct boundaries such as aco, dentate gyrus, granule cell layer (DG-sg), and CP were aligned well to the Allen reference atlas. For ACA, the left and right boundaries were not obvious, but the upper and lower boundaries could be judged by the original image. For MV and spinal nucleus of the trigeminal, interpolar part (SPVI) with inconspicuous boundaries, the accuracy could be roughly judged by the brain spatial orientation. All these areas were well aligned.

According to the above registration results, we first assessed the accuracy at brain-region level, a rough assessment (See method). We evaluated ten brain regions in brain-wide for the five datasets. The results are shown in box plots (Figure 7A). The median dice score of all brain regions and datasets were above 0.9.

**Figure 6.**
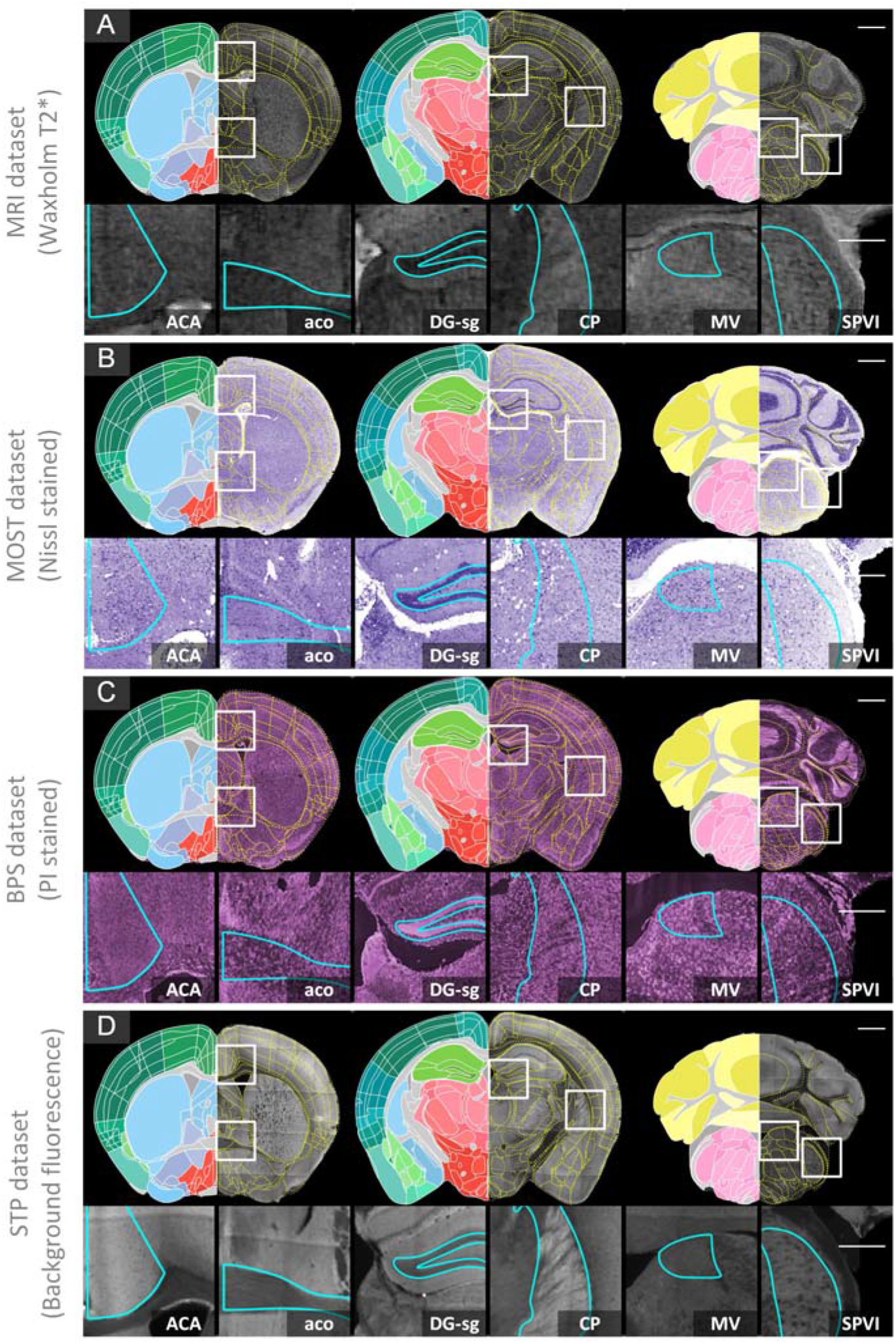
Registration effects for multi-modal datasets. The 10 μm thick projections of three coronal sections, the left half is the Allen CCFv3, and right half is the yellow dotted line of the Allen CCFv3 superimposed on original image. Enlarged views of several local regions are provided below indicated with white boxes in the above coronal sections. The projection thickness is 10 μm. (A): Dataset 2 (MRI), (B): Dataset 3 (MOST), (C): Dataset 4 (BPS), (D): Dataset 6 (STP). Scale bars: coronal 1 mm, detail 0.5 mm. Anterior cingulate area (ACA), anterior commissure, olfactory limb (aco), dentate gyrus, granule cell layer (DG-sg), caudoputamen (CP), medial vestibular nucleus (MV), spinal nucleus of the trigeminal, interpolar part (SPVI).

**Figure 7.**
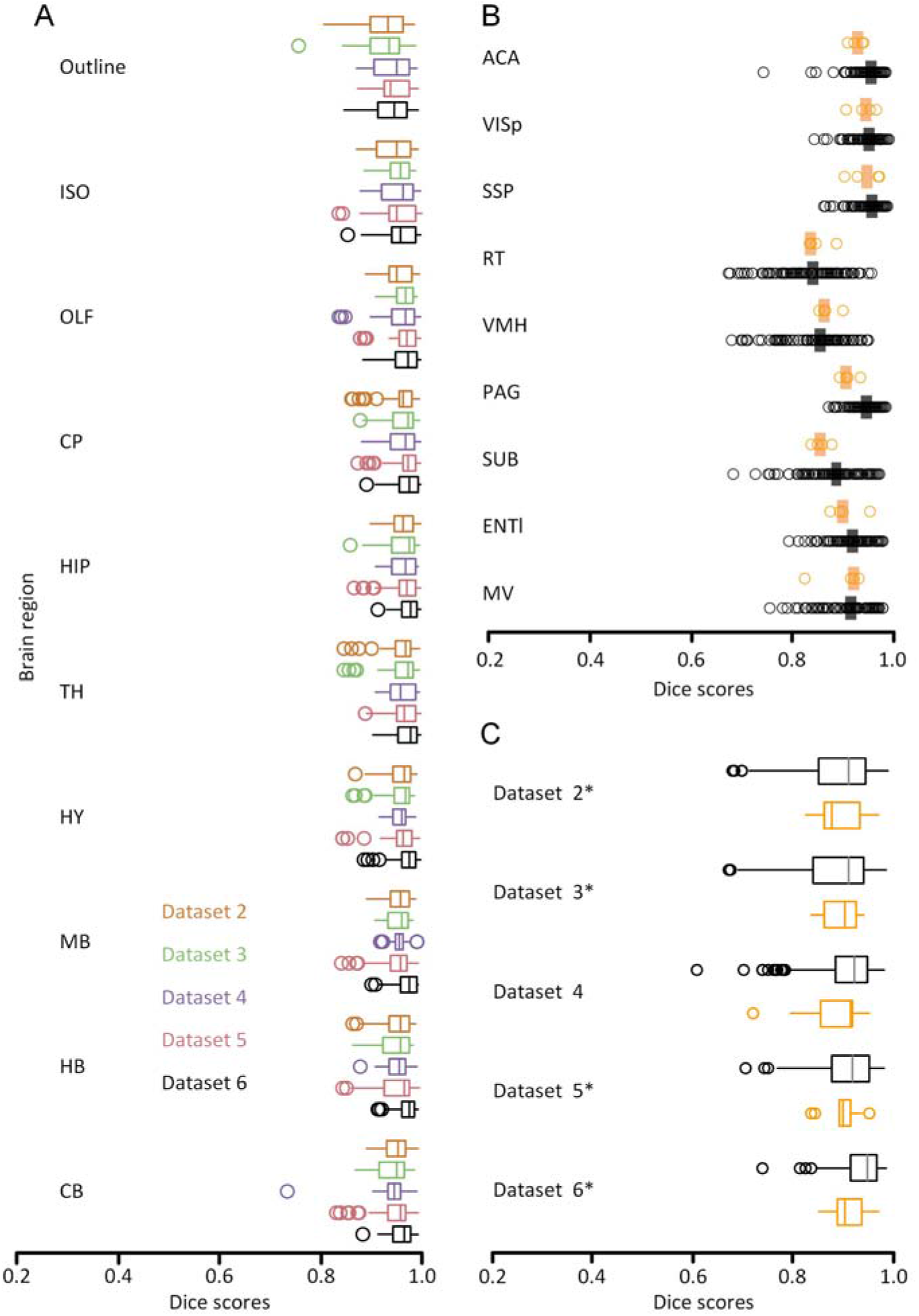
Multilevel quantitative assessment results. (A) The assessment at the brain-regions level, boxplot of dice scores for ten brain regions of five brain datasets (indicated with different colors) (n = 5). (B) The assessment at the nuclei level, dice scores of registration performance (RP, orange) and human performance (HP, black) are grouped by structure (n = 4). Vertical lines indicate the median scores. (C) The assessment at the nuclei level, dice scores of registration performance (RP, orange) and human performance (HP, black) are grouped by dataset (n = 5). Brains used in (B) are marked with an asterisk.

In addition to the assessment of the identifiable nuclei level at 10 μm resolution, nine brain anatomical structures were selected (see Method). When we grouped the results by HP or RP, the median acquired by HP (Figure 7B, black) was not significantly different from acquired by RP (Figure 7B, orange) (Mann-Whitney U-test, score of 0.90 versus 0.92, P = 0.054; n = 4 brains, 9 structures, 22 human raters). When we grouped these scores by nucleus, there were no significant differences between HP and RP in eight structures except the PAG. HP was significantly better than RP in the segmentation of PAG regions (Figure 7B) (Mann-Whitney U-test, score of 0.94 versus 0.90, P = 0.017; n = 4 brains, 9 structures, 22 human raters). When we grouped the results by datasets, there was also no significant difference between HP and RP in four individual brains (Figure 7B) (Mann-Whitney U-test, P > 0.06).

All these results demonstrated that the method proposed in this paper has extremely high accuracy at the brain-region level, and there was no significant difference from manual results at the nuclei level of 10 μm resolution.

### Whole-brain registration at single-cell resolution with TB-scale dataset

We used the low-resolution registration result (10 μm) to demonstrate the robustness and accuracy of BrainsMapi in the previous section. Moreover, we achieved nonlinear registration of TB-scale dataset at single-cell resolution.

Nonlinear registration at single-neuron resolution can not only correct non-uniform deformation of brain structures but also integrate various dataset at cellular level. Based on BrainsMapi (See Method), we aligned Dataset 5 (fluorescence channel) to Dataset 1 (Allen CCFv3) at single-cell resolution.

The Dataset 5 of 0.32 μm × 0.32 μm × 1 μm original resolution was sampled to 0.32 μm isotropic, and the dataset size reached 20 TB approximately. We mapped the three-dimensional, continuous, and single-cell resolution registration results to the standard atlas space. The dataset was aligned to the Allen CCFv3 in three-dimension (Figure 8ABC), and the brain regions were matched accurately (Figure 8G). We also presented the registration effects by comparing the pre- and post-registration of a coronal image with the Allen CCFv3 superimposed (Figure 8DG). The neuron fiber morphology was corrected and the complete and continuous structure was simultaneously achieved (Figure 8EH). Fine and weak signals also remained consistent after registration.

**Figure 8.**
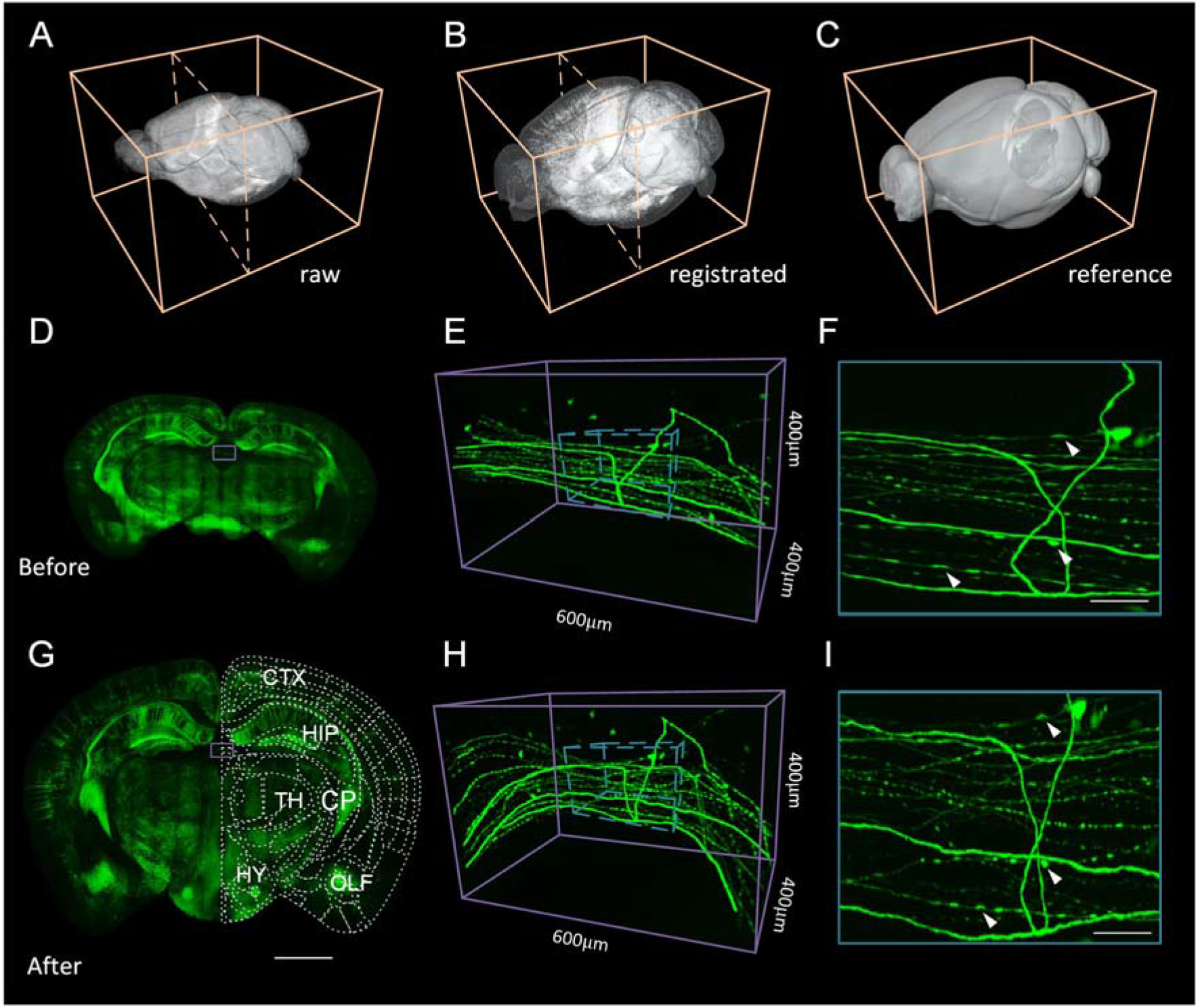
Registration effects for the whole-brain at single-cell resolution. (A) The three-dimensional rendering of the original GFP channel of Thy1-GFP M-line transgenic mice. (B) The three-dimensional rendering of the registered GFP channel. (C) The three-dimensional rendering of the brain outline of the Allen CCFv3. (D) The 256 μm thick coronal projection of the original image. (E) The enlarged three-dimensional rending view at the location indicated by the blue box in (D). (F) The enlarged 256 μm thick projection view at the location indicated by the blue cuboid in (E). (G) The 256 μm thick coronal projection of the registered image. (H) The enlarged three-dimensional rendered view at the location indicated by the blue box in (G). (I) The enlarged 256 m thick projection view at the location indicated by the blue cuboid in (H).

During the process, we completed the parallel computation of about 20 TB sized dataset in 70 hours, which used 100 threads in 20-node HPC and cost 56 GB memory for each node. In the addition, we evaluated the performance of registration for large volume dataset (SI Figure 6 and Table 2).

### Register existing metadata to a reference atlas

Metadata refers to digitized, vectorized information acquired from a raw image dataset, such as vascular structure, neuron projections, and cell distribution. These metadata are scattered in different brain spaces, individual laboratories and projects. Using BrainsMapi, vectorized dataset could be registered into a standard brain space to complete the integration of these existing and labeled metadata.

By aligning Metadata 1 to the Allen CCFv3, we presented the registration results of vectorized neurons. Vertebral neurons were manually traced from unregistered Brain 4 (Figure 9A). Using the proposed vectorized registration method, we deformed and completed the spatial localization of the metadata (Figure 9B). For comparison, we also presented the results of three-dimensional pre- and post-registration in horizontal, sagittal and coronal (Figure 9C).

**Figure 9.**
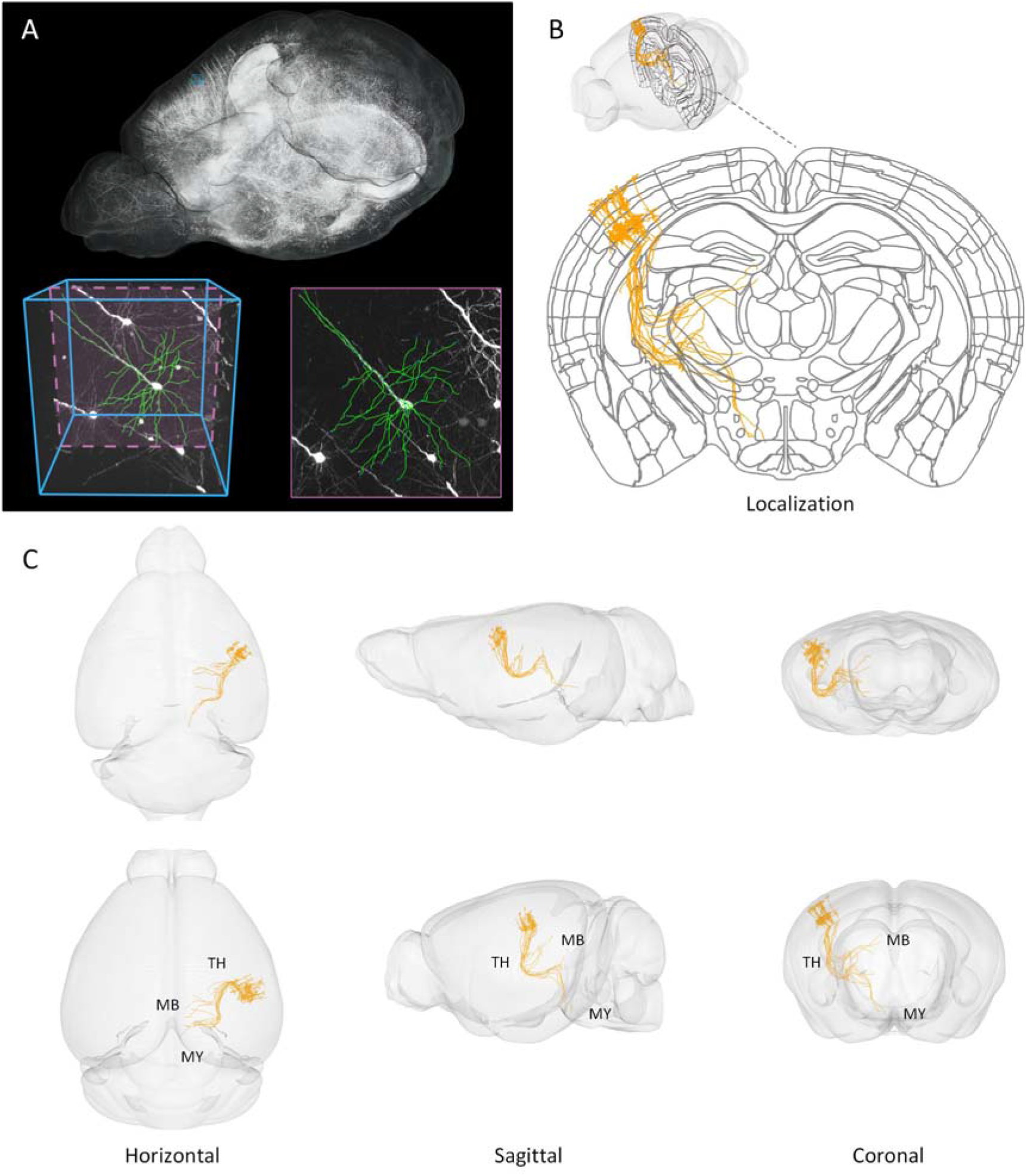
Registration effects for existing metadata. (A) The three-dimensional rendering of the GFP channel of Thy1-GFP M-line transgenic mice, three-dimensional and two-dimensional vectorized neurons (Metadata 1). (B) The coronal projection of the Allen CCFv3 with vectorized neurons after registration indicated in the three-dimensional rendering in (B). (C) The three-dimensional rendering of three anatomical projections (horizontal, sagittal and coronal) of the outline and neurons before (C top) and after registration (C bottom).

These results proved that the proposed method of registration of the vectorized dataset could complete the integration of metadata from different individuals.

## Discussion

In this study, we proposed a robust registration interface for large volume brain datasets named BrainsMapi. We used anatomically invariant regions during the registration to ensure the objective extraction of a large number of features, making it able to accurately register various datasets. Furthermore, with the low-resolution acquisition parameters and high-resolution transformation strategy, we realized the nonlinear registration of the TB-scale whole-brain dataset by data partitioning. We demonstrated the robustness of our registration method on both model data and real brain image datasets and presented the nonlinear registration results of a three-dimensional whole-brain fine image dataset at single-neuron resolution. Additionally, the labeled and existing vectorized datasets were registered to a standard brain space. Finally, we designed an objective multilevel evaluation method to prove the accuracy.

The regional features of our method are no limited to the cytoarchitectural image data, reflecting its wide applicability. As long as the anatomical region can be identified in the image, the method can be used for registration. However, the selection of regional features is not absolute, and can be autonomous based on the image characteristics or experimental requirements until the registration results meet expectations. An ideal situation is to select all the brain regions for registration, which will obtain the best results, but the cost is too high. In this paper, we recommended 14 regions based on our experience, aiming to obtain sufficiently accurate results with lower costs.

Another important aspect of this paper is the high-resolution nonlinear registration method for large volume datasets, which benefited from the low-resolution acquisition parameters, high-resolution transformation strategies, and data partitioning ideas; this method is also highly scalable and very promising for future applications to Peta Voxel datasets, such as the marmoset and human brain datasets. Presently, there appear to be no feasibility problems, and the only cost is more computing time, which can be accelerated by improving hardware performance.

Neuroscientific analysis with the brain spatial orientation requires matching the dataset to the standard brain space coordinate system to obtain anatomical boundaries. More general and common analysis skills involve combining multi-modality and multi-scale datasets to reveal the structural and functional relationship of the brain, such as MRI, optical imaging, even electron microscopy datasets. The integration of mesoscopic and macroscopic datasets is more meaningful, valuable and efficient. Many projects (Ascoli et al., 2007; Hawrylycz et al., 2011; Rosen et al., 2000) have used registration techniques to integrate various datasets even the recent international brain projects (Huang and Luo, 2015) desire to develop a powerful, standardized, industrialized framework to integrate multi-scale, multi-mode and massive datasets for studies on brain function mapping, disease models, and behavioral cognition (Okano et al., 2016). BrainsMapi is highly compatible with the requirements of data integration. It can accurately register various image datasets and existing vectorized metadata, and handle high-throughput handle the TB-scale large volume whole-brain datasets, providing a complete and effective pipeline for brain data integration.

## Acknowledgments

We appreciate X. Peng, W. Shi, and J. Yu for image preprocessing and Y. Li for TDat tools, and the financial supports from the 973 projection (Grant No.2015CB755602), Science Fund for Creative Research Group of China (Grant No.61721092), the director fund of the WNLO and the Foundation for the Author of National Excellent Doctoral Dissertation of PR China (FANEDD) (Grant No. 201464).

